# Infection, recovery and re-infection of farmed mink with SARS-CoV-2

**DOI:** 10.1101/2021.05.07.443055

**Authors:** Thomas Bruun Rasmussen, Jannik Fonager, Charlotte Sværke Jørgensen, Ria Lassaunière, Anne Sofie Hammer, Michelle Lauge Quaade, Anette Boklund, Louise Lohse, Bertel Strandbygaard, Morten Rasmussen, Thomas Yssing Michaelsen, Sten Mortensen, Anders Fomsgaard, Graham J. Belsham, Anette Bøtner

## Abstract

Mink, on a farm with about 15,000 animals, became infected with SARS-CoV-2. Over 75% of tested animals were positive for SARS-CoV-2 RNA in throat swabs and 100% of tested animals were seropositive. The virus responsible had a deletion of nucleotides encoding residues H69 and V70 within the spike protein gene. The infected mink recovered and after free-testing of the mink, the animals remained seropositive. During follow-up studies, after a period of more than 2 months without virus detection, over 75% of tested animals scored positive again for SARS-CoV-2 RNA. Whole genome sequencing showed that the virus circulating during this re-infection was most closely related to the virus identified in the first outbreak on this farm but additional sequence changes had occurred. Animals had much higher levels of anti-SARS-CoV-2 antibodies after re-infection than at free-testing. Thus, following recovery from an initial infection, seropositive mink rapidly became susceptible to re-infection by SARS-CoV-2.

**Article Summary Line:** Following widespread infection with SARS-CoV-2 of mink on a farm, all tested animals had seroconverted and the farm was then tested free of infection; however, less than 3 months later, a further round of infection affected more than 75% of tested animals.

## Introduction

The SARS-CoV-2 has caused a pandemic and contributed to the deaths of over 2 million people *(1)*. Farmed mink (*Neovison vison*) are also highly susceptible to infection by SARS-CoV-2 *(2, 3)*. As in humans, the infection in mink can cause respiratory distress and, in some cases, mortality. However, often the proportion of infected mink that show clinical disease is low. Cases of SARS-CoV-2 infection in farmed mink were initially observed in the Netherlands (NL), in April 2020 *(3)*, and then independently in Denmark (DK) in June 2020 (note, different clades of the virus were involved, see *(2)*). Outbreaks have continued and about 70 farms in the NL have been infected *(4)* while 290 farms out of about 1200 mink farms in DK were positive for the virus *(5)*. All mink (>15,000,000) have now been culled in DK *(6)*. Similarly, the termination of mink farming in the NL was brought forward by 3 years from the previously planned date of 1^st^ January 2024 *(4)*. The routes of transmission of the virus between mink farms are not fully understood *(5)* but it has become apparent that spread of the virus can occur not only from humans to mink but also from mink to humans *(2, 7)*.

After the initial cases of SARS-CoV-2 infection in mink in DK, on Farms 1-3 in Northern Jutland (as described in *(2)*), a regular screening program was established to test dead mink from all Danish mink farms for the presence of SARS-CoV-2, every 3^rd^ week *(6)*. Infection of mink on Farm 4 was identified through this Early Warning (EW) program but, in contrast to Farms 1-3, the mink were not culled and the seropositive animals apparently cleared the infection. This allowed an evaluation of the duration and efficacy of the immune response in mink to protect against re-infection.

## Results

### Infection of mink on Farm 4

Farm 4 (with about 2400 adult mink and 12600 kits housed in 24 open sheds), was located near Hjørring (also in Northern Jutland) and was tested as part of the EW screening program. On 20^th^ July 2020, as part of this system, 5 dead mink from this farm were tested for the virus and all were RT-qPCR negative. However, on 11^th^ August, a further 5 dead mink were tested and all were positive in this assay (Table 1). In follow-up testing, on 13^th^ August, 23 of 30 live mink tested (16 adults and 14 kits) were positive. A further 7 (of 10) dead mink also tested positive. All live mink tested (30 kits and 30 adults) were also strongly seropositive on 19^th^ August, but a reduced proportion of the mink (13 of these 60 mink tested) were positive for SARS-CoV-2 RNA. However, throat swab samples from 21 dead mink were all positive for viral RNA. Furthermore, on 31^st^ August, 7 out of 24 dead mink also tested positive by RT-qPCR. The mink on the farm were not culled but closely followed and, from 15^th^ September onwards, no virus was detected by RT-qPCR among the mink. For “ free-testing”, 300 animals were tested (in 60 pools of 5 samples) on 30^th^ September (Table 1) with negative results. This testing strategy was designed to detect, with 95% confidence, a 1% prevalence of SARS-CoV-2 RNA positive animals. Hence, the infection had apparently disappeared among the mink on this farm.

**Table 1.** Summary of laboratory analysis of mink sampling from Farm 4.

| Sample<br>origin | ELISA |  | RT-qPCR |  | Date of<br>sample collection |
| --- | --- | --- | --- | --- | --- |
|  | Sera<br>(positive/tested) | % | Throat swabs<br>(positive/tested) | % |  |
| Dead mink (EW) | n.d. |  | 0/5 | 0 | 20-07-2020 |
| Dead mink (EW) | n.d. |  | 5/5 | 100 | 11-08-2020<sup>1</sup> |
| Live adult mink | n.d. |  | 11/16 | 69 | 13-08-2020 |
| Live mink kits | n.d. |  | 12/14 | 86 | 13-08-2020 |
| Dead mink | n.d. |  | 7/10 | 70 | 13-08-2020 |
| Live adult mink | 30/30 | 100 | 4/30 | 13 | 19-08-2020 |
| Live mink kits | 30/30 | 100 | 9/30 | 30 | 19-08-2020 |
| Dead mink | n.d. |  | 21/21 | 100 | 19-08-2020 |
| Dead mink | n.d. |  | 7/24 | 29 | 31-08-2020 |
| Dead mink | n.d. |  | 0/31 | 0 | 15-09-2020 |
| Dead mink | n.d. |  | 0/25 | 0 | 28-09-2020 |
| Live mink | n.d. |  | 0/60* | 0 | 30-09-2020 |
| Live mink | 60/60 | 100 | 0/60 | 0 | 05-10-2020 |
| Dead mink (EW) | n.d. |  | 1/2** | 50 | 02-11-2020 |
| Dead mink (EW) | n.d. |  | 1/2** | 50 | 04-11-2020 |
| Live mink | 30/30 | 100 | 23/30 | 77 | 06-11-2020 |
| Dead mink | n.d. |  | 3/5 | 60 | 06-11-2020 |

n.d. : not done
1: Samples were received at SSI on this date.
\*300 animals were tested in pools of 5, i.e. in 60 assays.
\*\* two pools of 5 samples test

Surveillance of the farm continued and, in early October, 60 live mink were tested and were again all found negative by RT-qPCR but all these mink remained seropositive (Table 1). Thus, no animals tested positive by RT-qPCR in September and October but there was a very high (100%) prevalence of antibodies against anti-SARS-CoV-2. However, unexpectedly, 1 of 2 pools of 5 dead mink tested, as part of the continuing EW program, on both 2^nd^ and 4^th^ November were found to be positive by RT-qPCR (Table 1). Consequently, a further 30 animals were tested on 6^th^ November and 23 (77%) were found SARS-CoV-2 RNA positive while 100% of these animals remained seropositive, as observed one month previously. In addition, 3 out of 5 additional dead mink were found positive by RT-qPCR. The Ct values for 15 of the 23 samples that were found positive for viral RNA were below 30. No specific clinical signs of respiratory disease were apparent on the farm, however the farmer had noticed reduced feed intake and some cases of diarrhea in the mink.

Titration, in the ELISA, of the anti-SARS-CoV-2 antibodies in seropositive serum samples collected from August onwards showed that much higher levels of these antibodies were present in the mink in November, following the second round of infection than in August or October (see Figure 1A) although the seroprevalence in the mink had been high throughout. In August, the median antibody titre observed (from 16 animals tested) was 800 (with values ranging from 100 to 3200), with just a single animal having the highest titre. In early October (at free-testing), the titres in 15 sera tested were higher (ranging from 800 to 12800, with 4 of the sera having titres of ≥6400, the median titre was 3200). However, in November, following the reappearance of RT-qPCR positive animals, from 22 sera tested, 16 of them had titres ≥6400 and 12 had titres of 25600, see Figure 1A). Thus, it is clear that the anti-SARS-CoV-2 antibody levels, as measured by the ELISA, in the mink were greatly enhanced following the re-infection. It was apparent that in some individual animals no change in antibody levels were apparent in November, presumably because not all the animals had been re-infected.

**Figure 1.**
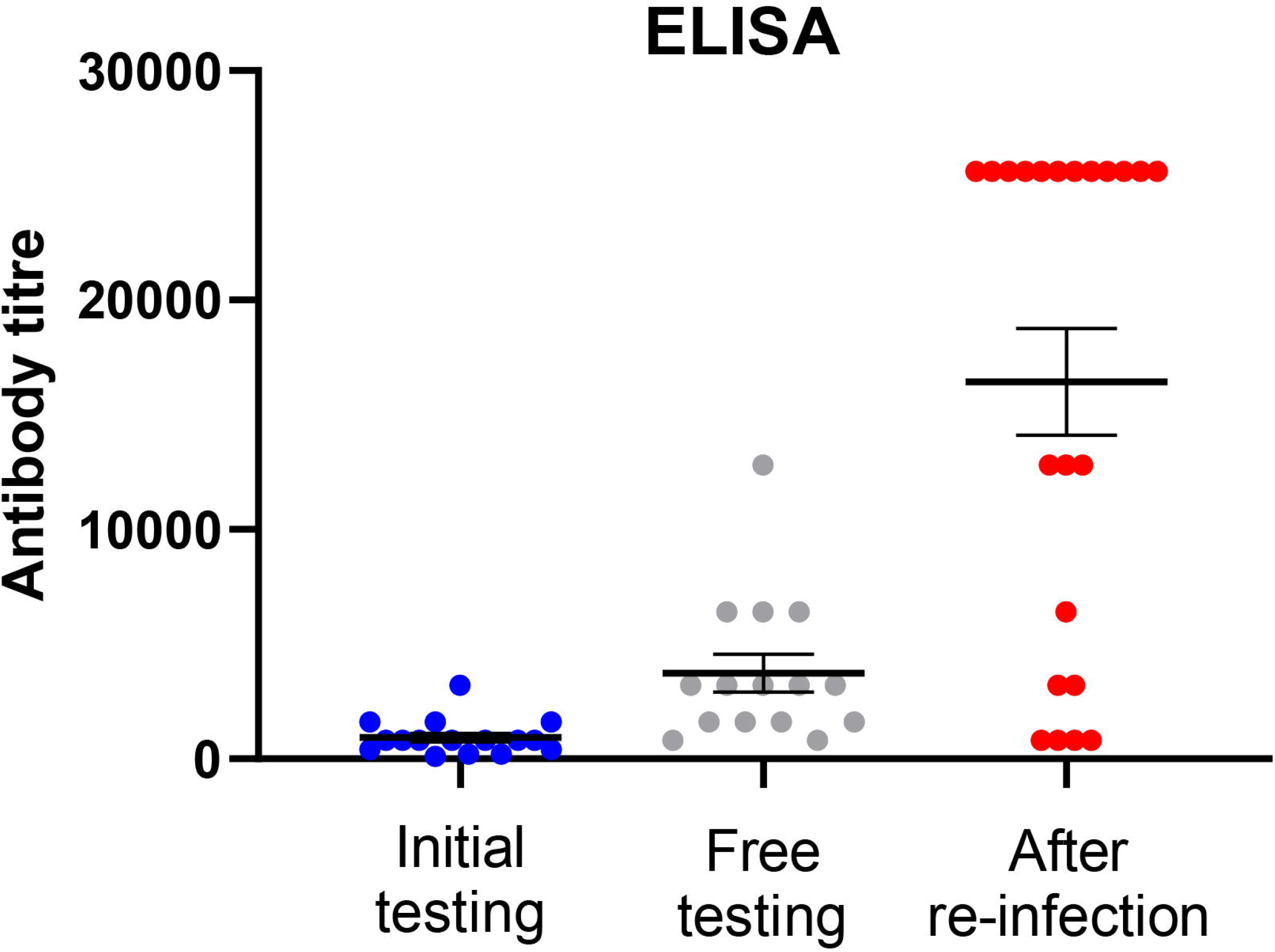
Panel A. Anti-SARS-CoV-2 antibody titres measured by ELISA. Selected positive sera from mink collected at the time of initial diagnosis (blue circles), at free-testing (grey circles) and following re-infection (red circles), on 19-08-20, 02-10-20 and 06-11-20 respectively, were titrated and assayed by ELISA. The reciprocals of the highest dilution yielding a positive signal are plotted. Mean (+/-SEM) values are indicated by horizontal black lines. Panel B. The same serum samples were also assayed in virus neutralization assays and the calculated antibody titres are plotted using the same colour scheme.

### Assessment of neutralizing antibodies

To assess whether the anti-SARS-CoV-2 antibodies in the mink that were detected by ELISA were capable of neutralizing virus infectivity, the same serum samples were also tested in virus neutralization assays using a human SARS-CoV-2 isolate that had the same amino acid changes in the spike protein as the viruses identified initially on Farm 4 (as used previously *(8)*). The results (see Figure 1B) showed a similar pattern of antibody development as observed in the ELISA. All the ELISA positive sera tested had neutralization activity but the levels of these antibodies were greatly elevated after the second round of infection (sera collected in November). There was a high degree of correspondence between the levels of antibodies detected in the two different types of assay (for all samples, the Spearman correlation co-efficient r= 0.793, p <0.0001).

### Whole genome sequencing of viruses on Farm 4

The complete genome sequences of the viruses, from multiple samples, from infected mink on Farm 4 in August and in November were determined. The viruses present on Farm 4 in August were all from clade 20B and were very closely related to the viruses that were identified on Farms 1, 2 and 3 *(2)* (Table 2 and Figure 2) and appeared to be part of the same transmission chain. In particular, they each had the mutation A22920T in the spike protein coding sequence, resulting in the amino acid substitution Y453F, which is a hallmark of most viruses that have infected mink in DK. This change was also detected on farm NB02, one of four farms initially infected in NL, however, this was within a different virus clade (see *(2, 3)*). However, in addition to this change, the spike protein gene in the viruses on Farm 4 also had a deletion of 6 nt (Δ21766-21771). This deletion affects 3 separate codons, changing GCT.ATA.CAT.GTC.TCT to GCT.ATC.TCT, the encoded amino acid sequence is changed from A-I-H^69^-V^70^-S to A-I-S thus residues H69 and V70 in the N-terminal domain (NTD) of the spike protein are lost. This deletion had not been identified previously in mink or in humans in combination with the Y453F substitution (see Table 2) but the deletion of these residues is shared with the SARS CoV-2 variant of concern (VOC) 202012/01 *(9)*. Two other deletions in the ORF1a coding sequence (Δ517-519 and Δ6510-6512) and two other amino acid substitutions (P3395S in ORF1a and S2430I in ORF1b) were also observed in some of the viruses present in the mink during this initial infection in August. The viruses present on Farm 4 in November were most closely related to those seen previously on Farm 4, over 2 months earlier (Figure 2). It should be noted that, by November 2020, over 200 farms in DK had been identified as having infected mink *(5)* and a number of different variants had been observed in the animals *(6)*. The viruses on Farms 1-3 were closely related to each other and also to the viruses present in August on Farm 4, but the latter viruses had some additional changes (e.g. the deletion of residues H69 and V70 in the spike protein, see Table 2), which persisted throughout the rest of the outbreaks in farmed mink. Thus, viruses in farms infected after Farm 4 (identified on August 11^th^) were nearly all derived from those first detected on Farm 4. As indicated above, the November viruses from Farm 4 had the A22920T mutation and the deletions in the S and ORF1a coding sequences. However, the November viruses had additional changes across the genome, both within and outside of the S gene, compared to the viruses in Farms 2 and 3 (Table 2). It is noteworthy that the Farm 4 sequences in November had changes at nt 10448 (encoding the substitution P3395S in ORF1a) and 20756 (encoding S2430I in ORF1b) that had only been seen in a subset of the August sequences from Farm 4 (samples Farm4_18_13-08-2020 and Farm4_19_13-08-2020, see Figure 2). These changes act as a fingerprint and strongly suggest that it was not an entirely new introduction of virus into the farm from elsewhere. Furthermore, the viruses on Farm 4 in November also all shared changes at nt 3792 (resulting in A1176V), 5167, 10887 (resulting in G3541E), 21727 and 23815 (the latter two silent changes are in the S gene) that were not present in any of the Farm 4 sequences in August (Table 2). The presence of these additional sequence changes indicates that the virus had been replicating in hosts with close connection to this farm between August and November but does not prove that the virus has continued to replicate in mink during this time.

**Table 2.** Sequence changes within SARS-CoV-2 in mink on Farm 4.

| Location | 5'-UTR | ORF1a |  |  |  | ORF1b |  |  | S |  |  | ORF3a | N |  |
| --- | --- | --- | --- | --- | --- | --- | --- | --- | --- | --- | --- | --- | --- | --- |
| Nt | 241 | 3037 | 5144 | 10448 | 11776 | 14408 | 15656 | 20756 | Δ21766-21771 | 22920 | 23403 | 25936 | 28854 | other |
| Virus |  |  |  |  |  |  |  |  |  |  |  |  |  |  |
| Wuhan | C | C | C | C | C | C | C | G | - | A | A | C | C |  |
| EPI_ISL_455326 20B | T | T | C | C | C | T | C | G | - | A | G | C | C |  |
| Farm 1 | T | T | C | C | C | T | T | G | - | T/A | G | T | C |  |
| Farm 2 | T | T | C | C | C | T | T | G | - | T | G | T | C |  |
| Farm 3 | T | T | C | C | C | T | T | G | - | T | G | T | C |  |
| **Aug 2020** |  |  |  |  |  |  |  |  |  |  |  |  |  |  |
| Farm4_5 | T | T | T | C | T | T | T | G | + | T | G | T | T |  |
| Farm4_6 | T | T | T | C | T | T | T | G | + | T | G | T | T | G488A |
| Farm4_8 | T | T | T | C | T | T | T | G | + | T | G | T | T | Δ21984-21995 |
| Farm4_18 | T | T | T | T | T | T | T | T | + | T | G | T | T |  |
| Farm4_19 | T | T | T | T | T | T | T | T | + | T | G | T | T |  |
| Farm4_21 | T | T | T | C | T | T | T | G | + | T | G | T | T | A652C (K129N)<sup>1</sup> |
| Farm4_35 | T | T | T | C | T | T | T | G | + | T | G | T | T | Δ27982-28030 |
| Farm4_37 | T | T | T | C | T | T | T | G | + | T | G | T | T | T1873C,G2035T(L590F) |
| **Nov 2020** |  |  |  |  |  |  |  |  |  |  |  |  |  |  |
| Farm4_1 | T | T | T | T | T | T | T | T | + | T | G | T | T | C1913T (R550C), C3792T (A1176V), C5167T, G10887A (G3541E), C21727T, T23815C |
| Farm4_14 | T | T | T | T | T | T | T | T | + | T | G | T | T | A3303G, C3792T (A1176V), C5167T, G10887A (G3541E), C21727T, T23815C |
| Farm4_15 | T | T | T | T | T | T | T | T | + | T | G | T | T | A3303G, C3792T (A1176V), C5167T, G10887A (G3541E), C21727T, T23815C |
| AA change | - | - | - | P3395S | - | P314L | T730I | S2430I | ΔH69-V70 | Y453F | D614G | H182Y | S194L |  |

1: Note the same additional sequence change was also present in 4 other samples (Farm4\_16\_13-08-2020, Farm4\_20\_13-08-2020, Farm4\_22\_13-08-2020 and Farm4\_4\_19-08-2020). N.B. All the mink viruses, together with the EPI\_ISL455326 clade 20B representative sequence, shown here were from clade 20B and had G28881A, G28882A and G28883C changes compared to the Wuhan strain. In addition, the mink viruses from Farm 4 also lacked nt 517-519 and nt 6510-6512. Other nt changes from the Wuhan reference sequence are highlighted in yellow while nt changes from the representative clade 20B virus are shown in red type. Shared additional changes that occurred in viruses on Farm 4 between August and November 2020 are indicated with colour codes, encoded amino acid changes, where applicable, are shown in parenthesis.

**Figure 2.**
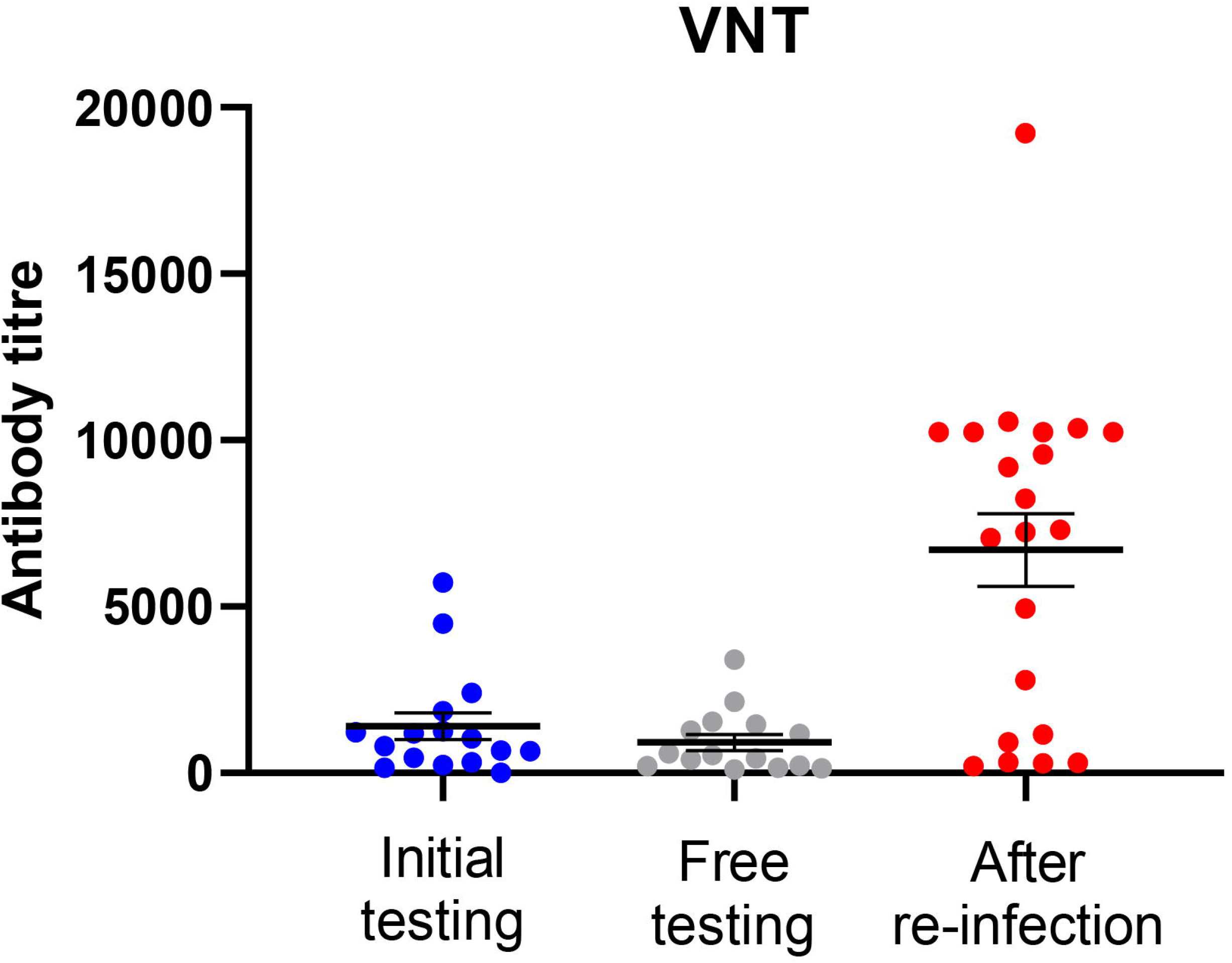
Phylogenetic tree showing the relationships between the full genome sequences of SARS-CoV-2 samples from Farms 1-4. Sequences from the re-infection (collected in November) are indicated in red while samples collected in August are indicated in blue.

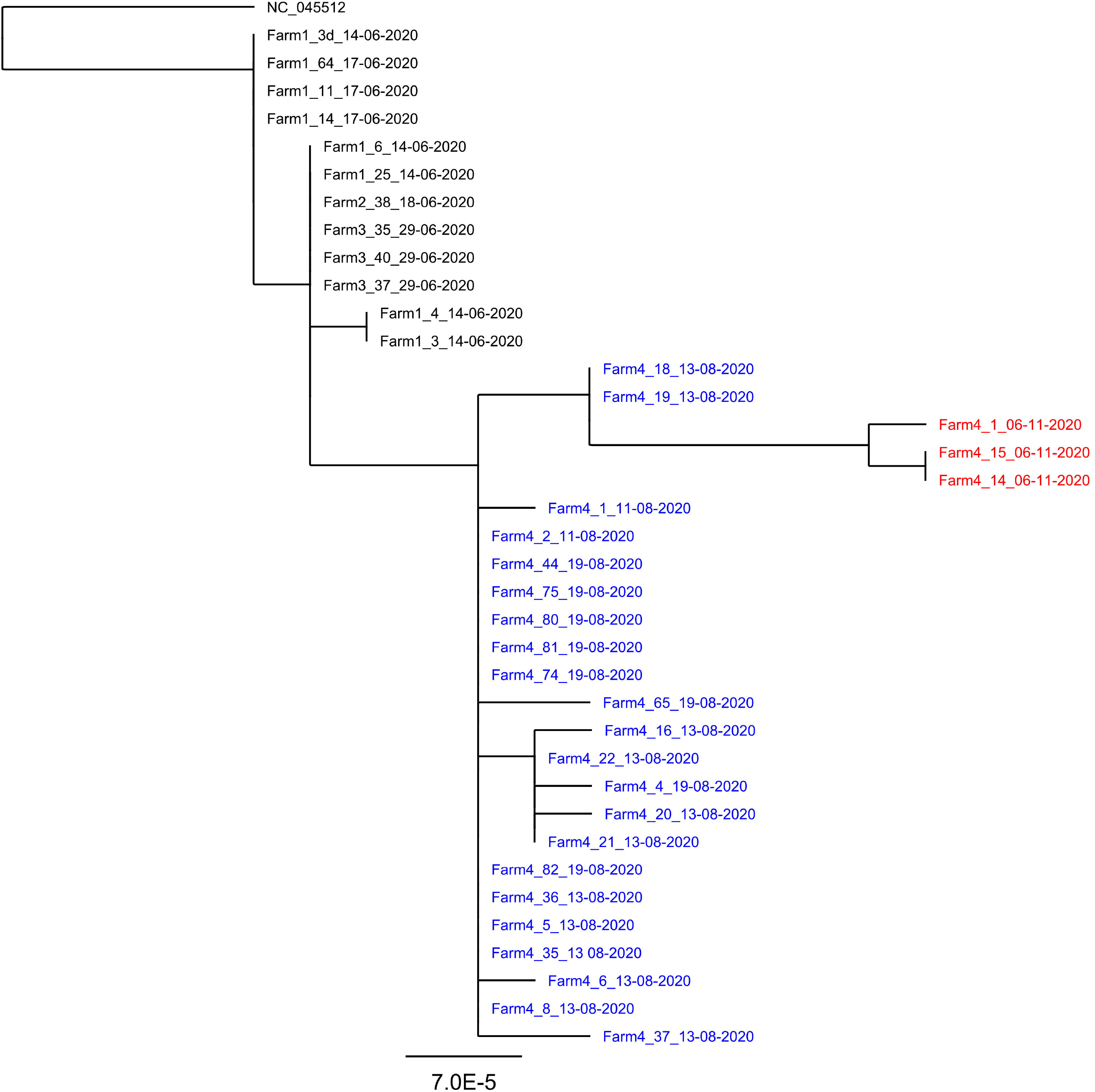

Phylogenetic analysis clearly showed that all viruses from Farm 4 were very closely related to each other, including the viruses from both August and November (Figure 2). As described above, two of the early Farm 4 viruses (Farm4_18_13-08-2020 and Farm4_19_13-08-2020) shared additional changes at nt 10448 and 20756 (see Table 2) and the November viruses formed their own distinct branch from these (Figure 2), due to the presence of the further sequence changes (Table 2).

## Discussion

SARS-CoV-2 can readily infect humans and mink. In addition, certain other species, e.g. cats, dogs and ferrets, can also be infected following direct inoculation under experimental conditions *(10, 11)*. Furthermore, some cases of transmission from infected people to their cats and dogs have occurred but it does not seem to happen more generally. Both cellular and humoral immune responses occur within SARS-CoV-2-infected people and animals *(12, 13)* and it is common for both humans and animals to be both seropositive and RT-qPCR positive simultaneously (see *(2, 14)*). However, as people and animals recover, the levels of virus subside but antibody levels persist, or increase, at least for some time.

Farm 4 was the only Danish mink farm, where the animals were allowed to recover and were tested with the purpose of documenting freedom from SARS-CoV-2 infection. Thus, Farm 4 gave a unique opportunity to follow the maintenance of anti-SARS-CoV-2 antibodies over an extended period and the resistance of the animals to reinfection. As observed on other mink farms in DK *(5)*, very widespread infection of the mink on Farm 4 by SARS-CoV-2 occurred in the first wave of infection, with 100 % of the tested animals being seropositive. As indicated above, the mink on Farm 4 were not culled after the detection of infection in August but during the following period of over 2 months, the animals were repeatedly screened and found to be negative by RT-qPCR, while the 100% seroprevalence remained. However, in November, it was observed that the mink had become infected again. A high proportion (>75%) of the animals tested had been re-infected by SARS-CoV-2 (Table 1). The virus responsible for the second round of infection was most closely related to the virus found almost 3 months earlier on this farm, with distinctive differences from the viruses responsible for the initial infections observed in mink on Farms 1-3 *(2)*. Notably, specific deletions were present within the spike protein gene and within the ORF1a gene in the virus responsible for the initial infection in August and in the later re-infection (Table 2). The virus acquired additional sequence changes during the period between the infections recognized in August and November, indicative of continued replication, rather than simply having been preserved in an infectious form. Since the virus present on Farm 4 in November was most closely related to virus present on the same farm in August, it seems most likely that re-infection of the mink from within the farm had occurred. It cannot be established, however, whether the virus had continued to replicate in a small number of mink on the farm, but with very restricted spread, or if it had replicated in an alternative host, linked to the farm, during this time and had then been re-introduced into the seropositive mink. It has been demonstrated, in both DK and the NL, that transmission between humans and mink can occur in both directions *(2, 6)*. Transmission to, and from, other hosts is theoretically possible (but not described previously; some cats were found to be infected on mink farms in DK and in the NL but they do not seem to spread the virus). It has been found that there was a cluster of occurrences of SARS-CoV-2 with the ΔH69/N70 and Y453F changes (as in Farm 4) in the local human population in August. Furthermore, a virus containing these changes plus the additional mutations (i.e. C3792T, C5167T, G10887A, C21727T and T23815C, see Table 2), which were present in the mink viruses from Farm 4 in November, was found in one person in the first half of November (data not shown). It seems likely that these human cases were infections derived from the mink.

A high proportion of the sequence changes observed in mink (see Table 2), which occurred in the viruses from Farm 4 between August and November (and also between the clade 20B viruses and the Wuhan virus, see *(2)*), involved C to T changes (in cDNA) that correspond to C to U changes in the viral RNA. Several of these nt changes are synonymous, i.e., they do not result in amino acid sequence changes. It has been suggested that such changes reflect host immune pressure via RNA editing systems (e.g. by APOBEC) rather than selection for increased transmissibility in particular hosts *(15, 16, 17)*. However, this process of RNA editing is not relevant to the key mutation in the S gene (A22920T), which seems to be an adaptation that occurred during the initial infection of mink *(2)*, or to the generation of deletions. The loss of residues H69 and V70 in the spike protein, seen in mink for the first time on Farm 4, and in certain variants from people, has been reported to double the infectivity of pseudoviruses displaying the mutant spike protein compared to the wild type particles *(18)*.

The sampling of the mink on Farm 4 tested, at most, 300 animals on any particular date, out of a population of about 15,000 animals. The free-testing strategy was designed to detect 1% prevalence with high (95%) confidence. It is clearly possible that a small number of infected animals were missed although the repeated follow-up screening makes this unlikely. However, the level of seroprevalence prior to the second round of infection had remained very high (100%) in the animals tested. Thus, it is not clear why so many animals (77% of 30 animals tested) were susceptible to a second round of infection. It has been considered whether the seropositivity detected in kits in August may be a consequence of maternally derived antibodies that could potentially decline more rapidly than antibodies generated from the infection in each animal. However, it seems difficult to reconcile this with the fact that >80% of throat swabs from mink kits tested clearly positive by RT-qPCR in August, which indicated a high level of infection amongst the kits in the first wave also.

The measurements of antibody responses were made using an ELISA that targets the receptor binding domain (RBD) of the spike protein. Antibodies to the SARS-CoV-2 spike protein were present in up to 100% of the infected mink. The antibody titres, measured in this assay, increased to very high levels during the period of re-infection (see Figure 1A). In studies on human sera, samples testing clearly positive (10 x cut-off) in this ELISA all had neutralizing antibodies *(19)*. Indeed, assessment of the same samples of mink sera as tested by ELISA in virus neutralization tests indicated a high correspondence between these two types of assay. Thus, the ELISA positive mink sera neutralized the virus and, furthermore, the sera collected in November, after reinfection, had much higher levels of anti-SARS-CoV-2 antibodies as measured in each assay (Figure 1B).

It appears that the virus responsible for the infections in November was not antigenically distinct from the virus in August since there were no non-synonymous changes within the spike protein gene during this time, although some silent sequence changes (i.e. C21727T and T23815C) had occurred, as well as changes elsewhere, within the virus genome, as usually occurs.

The most plausible conclusion is that infection of farmed mink with SARS-CoV-2 does not induce long-term protection against the virus. This should be compared with the situation in rhesus macaques where primary infection did protect against reinfection at about 1 month post-initial infection *(13, 20)* and in humans where protection from reinfection may last at least eight months *(12, 21)*. However, some cases of re-infection have been reported in health care workers in Brazil *(22)*, although this seems to have occurred in people who only developed a weak immune response during the initial infection. Furthermore, only about 50% of people in DK, who were over 65 years of age and had been infected with SARS-CoV-2, were found to be protected against re-infection *(23)*. On a mink farm, a large number of animals live in close proximity to each other and, potentially, once the infection occurs in some animals then there can be a rapid increase in virus production and a strong challenge to neighbouring animals. Perhaps this is sufficient to overcome the immune response. It is notable that greatly enhanced levels of anti-SARS-CoV-2 antibodies were detected in the mink following the second round of infection (Figure 1), but this was also observed following challenge of previously infected rhesus macaques which did not become re-infected *(13, 20)*. Currently, there are no “ correlates of protection” that can be used to evaluate the immune responses in mink.

## Methods

Blood and throat-swab samples were collected from mink (adults and kits) as indicated in Table 1. The presence of SARS-CoV-2 RNA was determined by RT-qPCR *(2)*. The SARS-CoV-2 Ab ELISA (Beijing Wantai Biological Pharmacy Enterprise, Beijing, China) was performed as described by the manufacturer, with the addition of an extra titration of positive samples.

Antibody titres are presented as the reciprocal of the highest dilution of the serum giving a positive result. Neutralising antibody titres were determined as described previously *(8)*. SARS-CoV-2 positive RNA samples were sequenced as described *(2)* and SARS-CoV-2 sequences were aligned using MAFFT *(24)*. Phylogenetic analysis was performed using the Maximum Likelihood method with the General-Time-Reversible model *(25)*.

## Acknowledgements

We greatly appreciate the collaborative attitude of the farmer. Furthermore, we appreciate the work done by technicians and official veterinarians at the Danish Veterinary and Food Administration (FVST) who collected many of the samples. The Danish COVID-19 Genome Consortium (https://www.covid19genomics.dk) is acknowledged for sequencing.

## Author Bio

Dr. Thomas Bruun Rasmussen, a senior researcher at the Statens Serum Institut in Copenhagen, is a veterinary virologist with special interest and expertise in molecular methods applied to diagnostics, research, and surveillance of transboundary animal diseases.

## References

1. Johns Hopkins University coronavirus resource center, https://coronavirus.jhu.edu/

2. Hammer AS, Quaade ML, Rasmussen TB, Fonager J, Rasmussen M, Mundbjerg K, et al. SARS-CoV-2 Transmission between Mink (Neovison vison) and Humans, Denmark. Emerg Infect Dis. 2021;27(2):547–51. doi: 10.3201/eid2702.203794.

3. Oreshkova N, Molenaar RJ, Vreman S, Harders F, Munnink BBO, Hakze-van der Honing RW et al. SARS-CoV-2 infection in farmed minks, the Netherlands, April and May 2020. Euro Surveill. 2020;25:2001005. https://doi.org/10.2807/1560-7917.ES.2020.25.23.2001005

4. https://promedmail.org/promed-posts/id=20201204.7994061

5. Boklund A, Hammer AS, Quaade ML, Rasmussen TB, Lohse L, Strandbygaard B, et al. SARS-CoV-2 in Danish Mink Farms: Course of the Epidemic and a Descriptive Analysis of the Outbreaks in 2020. Animals (Basel). 2021;11(1):164. doi: 10.3390/ani11010164.

6. Larsen HD, Fonager J, Lomholt FK, Dalby T, Benedetti G, Kristensen B, et al. Preliminary report of an outbreak of SARS-CoV-2 in mink and mink farmers associated with community spread, Denmark, June-November 2020. Euro Surveill. 2021;26(5). doi: 10.2807/1560-7917.ES.2021.26.5.210009

7. Oude Munnink BB, Sikkema RS, Nieuwenhuijse DF, Molenaar RJ, Munger E, Molenkamp R, et al. Transmission of SARS-CoV-2 on mink farms between humans and mink and back to humans. Science. 2020 Nov 10:eabe5901. doi: 10.1126/science.abe5901.

8. Lassaunière, R, Fonager, J, Rasmussen, M, Frische, A, Polacek, C, Rasmussen, TB, et al. In vitro characterization of fitness and convalescent antibody neutralisation of SARS-CoV-2 Cluster 5 variant emerging in mink at Danish farms. (submitted).

9. Bal A, Destras G, Gaymard A, Stefic K, Marlet J, Eymieux S, et al. Two-step strategy for the identification of SARS-CoV-2 variant of concern 202012/01 and other variants with spike deletion H69-V70, France, August to December 2020. Euro Surveill. 2021 Jan;26(3):2100008. doi: 10.2807/1560-7917.ES.2021.26.3.2100008.

10. Shi J, Wen Z, Zhong G, Yang H, Wang C, Huang B, et al. Susceptibility of ferrets, cats, dogs, and other domesticated animals to SARS–coronavirus 2. Science. 2020;368:1016–20. https://doi.org/10.1126/science.abb7015

11. Schlottau K, Rissmann M, Graaf A, Schön J, Sehl J, Wylezich C, et al. SARS-CoV-2 in fruit bats, ferrets, pigs, and chickens: an experimental transmission study. The Lancet Microbe. 2020; 1(5):e218–25. https://doi.org/10.1016/S2666-5247(20)30089-6

12. Hartley GE, Edwards ESJ, Aui PM, Varese N, Stojanovic S, McMahon J, et al.. Rapid generation of durable B cell memory to SARS-CoV-2 spike and nucleocapsid proteins in COVID-19 and convalescence. Sci Immunol. 2020;5(54):eabf8891. doi: 10.1126/sciimmunol.abf8891.

13. Chandrashekar A, Liu J, Martinot AJ, McMahan K, Mercado NB, Peter L, et al. SARS-CoV-2 infection protects against rechallenge in rhesus macaques. Science. 2020 369(6505):812–17. doi: 10.1126/science.abc4776.

14. Hung IF, Cheng VC, Li X, Tam AR, Hung DL, Chiu KH, et al. SARS-CoV-2 shedding and seroconversion among passengers quarantined after disembarking a cruise ship: a case series. Lancet Infect Dis. 2020;20(9):1051–60. doi: 10.1016/S1473-3099(20)30364-9.

15. Simmonds P. Rampant C?U hypermutation in the genomes of SARS-CoV-2 and other coronaviruses: causes and consequences for their short-and long-term evolutionary trajectories. mSphere 5, e00408–20 (2020).

16. Di Giorgio S, Martignano F, Torcia MG, Mattiuz G, Conticello SG. Evidence for host-dependent RNA editing in the transcriptome of SARS-CoV-2. Sci Adv 2020. doi:10.1126/sciadv.abb5813.

17. van Dorp L, Richard D, Tan CCS, Shaw LP, Acman M, Balloux F. No evidence for increased transmissibility from recurrent mutations in SARS-CoV-2. Nat Commun 11, 5986 (2020). https://doi.org/10.1038/s41467-020-19818-2

18. Kemp SA, Collier DA, Datir RP, Ferreira IATM, Gayed S, Jahun A, et al., SARS-CoV-2 evolution during treatment of chronic infection. Nature. 2021 Feb 5. doi: 10.1038/s41586-021-03291-y.

19. GeurtsvanKessel, CH, Okba N, Igloi Z, Bogers S, Embregts C, Laksono BM, et al. An evaluation of COVID-19 serological assays informs future diagnostics and exposure assessment. Nature Communications. 2020; 11(1), 3436. https://doi.org/10.1038/s41467-020-17317-y

20. Deng W, Bao L, Liu J, Xiao C, Liu J, Xue J, et al. Primary exposure to SARS-CoV-2 protects against reinfection in rhesus macaques. Science. 2020; 369(6505):818–823. doi: 10.1126/science.abc5343.

21. Choe PG, Kim K-H, Kang CK, Suh HJ, Kang E, Lee SY, et al. Antibody responses 8 months after asymptomatic or mild SARS-CoV-2 infection. Emerg Infect Dis. 2021. Epub https://doi.org/10.3201/eid2703.204543

22. Adrielle Dos Santos L, Filho PGG, Silva AMF, Santos JVG, Santos DS, Aquino MM et al. Recurrent COVID-19 including evidence of reinfection and enhanced severity in thirty Brazilian healthcare workers. J Infect. 2021:S0163-4453(21)00043-8. doi: 10.1016/j.jinf.2021.01.020.

23. Hansen CH, Michlmayr D, Gubbels SM, Mølbak K, Ethelberg S. Assessment of protection against reinfection with SARS-CoV-2 among 4 million PCR-tested individuals in Denmark in 2020: a population-level observational study. Lancet. 2021:S0140-6736(21)00575-4. doi: 10.1016/S0140-6736(21)00575-4

24. Katoh K, Standley DM. MAFFT Multiple Sequence Alignment Software Version 7: Improvements in Performance and Usability. Molecular Biology and Evolution. 2013; 30(4):772–780. doi:10.1093/molbev/mst010

25. Guindon S, Dufayard JF, Lefort V, Anisimova M, Hordijk W, Gascuel O. New Algorithms and Methods to Estimate Maximum-Likelihood Phylogenies: Assessing the Performance of PhyML 3.0, Systematic Biology. 2010; 59(3):307–321, doi:10.1093/sysbio/syq010

